# Limited response of primary nasal epithelial cells to *Bordetella pertussis* infection and the effector protein BteA

**DOI:** 10.1101/2025.02.03.636266

**Authors:** Martin Zmuda, Barbora Pravdova, Ivana Malcova, Ondrej Cerny, Denisa Vondrova, Jana Kamanova

## Abstract

*Bordetella pertussis* is a Gram-negative coccobacillus that causes whooping cough or pertussis, a respiratory disease that has recently experienced a resurgence. Upon entering the respiratory tract, *B. pertussis* colonizes the airway epithelium and attaches to ciliated cells. Here, we used primary human nasal epithelial cells (hNECs) cultured at the air-liquid interface, and investigated their interaction with the *B. pertussis* B1917, focusing on the role of the type III secretion system effector protein BteA. In this model, which resembles the epithelial cells of nasal epithelium *in vivo*, *B. pertussis* B1917 initially replicated in the overlying mucus and scarcely colonized the cell cilia. The colonization led to a gradual decline in epithelial barrier function, as shown by measurements of trans-epithelial electrical resistance (TEER) and staining of the tight junction protein zonula occludens 1 (ZO-1). The decrease in TEER occurred independently of the cytotoxic effector protein BteA. Transcriptomic and proteomic analyses of hNECs showed only moderate changes following infection, primarily characterized by increased mucus production, including upregulation of mucin MUC5AC. No profound response to BteA was detected. Furthermore, the infection did not induce production of inflammatory cytokines, suggesting that *B. pertussis* B1917 evades recognition by hNECs in this model system. These findings suggest that the bacterium may utilize the mucus layer in the airways as a protective niche to minimize epithelial recognition and damage.

**Importance:** The nasal epithelium is the initial site where *B. pertussis* comes into contact with the host during respiratory infection. This work established human nasal epithelial cells (hNECs) cultured at the air-liquid interface (ALI) as an *in vitro* model to investigate *B. pertussis* infection. Using this system, we were able to show that in the early stages of colonization the clinical isolate *B. pertussis* B1917 replicates in the mucus without disrupting epithelial barrier function. Infection results in moderate transcriptomic and proteomic changes and is characterized by increased mucus production and minimal inflammatory signaling.

## Introduction

*Bordetella pertussis*, a Gram-negative coccobacillus, is the causative agent of whooping cough, or pertussis. This highly transmissible respiratory disease presents as an acute, often fatal illness in infants and as a persistent, severe paroxysmal cough with characteristic whooping sounds in adults (1). Pertussis continues to be a significant health concern and it was estimated that in 2014 there were 24 million cases in children under 5 years with 160,000 deaths (2). While the highest incidence is observed in unvaccinated populations, a resurgence of the disease has been seen in developed countries that have transitioned from whole-cell pertussis (wP) vaccines to less reactive acellular pertussis (aP) vaccines (3). Current aP vaccines have been demonstrated to be insufficient in preventing nasal colonization by *B. pertussis* (4). This is likely due to their restricted ability to elicit strong mucosal immunity, including induction of secretory IgA as well as IL-17 and IFN-γ-producing respiratory tissue-resident memory T cells (T_RM_) (5–8). As a result, subclinical infections with *B. pertussis* are relatively common in aP-vaccinated children and adolescents, which poses a considerable risk of transmission to unvaccinated or incompletely vaccinated infants (9). Recently, several developed countries, including the Czech Republic, have reported outbreaks of pertussis. This resurgence may be related to the reduced circulation of *B. pertussis* during the COVID-19 pandemic, which led to increased susceptibility of the population (10).

*B. pertussis* is a pathogen exclusively adapted to humans with no animal or environmental reservoirs, which is transmitted via respiratory droplets. After inhalation, *B. pertussis* has been reported to attach to the ciliated cells and colonize the ciliated pseudostratified columnar epithelium of the respiratory tract (11, 12). This epithelium consists of ciliated cells and secretory club and goblet cells, which are essential for mucociliary clearance and barrier function. Besides, multipotent basal cells are responsible for epithelial regeneration (13, 14). However, advances in single-cell transcriptomics have further identified rarer cell types within the airway epithelium, including tuft cells, pulmonary neuroendocrine cells and ionocytes, and various transitional and intermediate cell types (13–16). The airway epithelium also actively contributes to respiratory defense by secreting antimicrobial substances and interacting with stromal, neuronal, and immune cells (both tissue-resident and recruited), to maintain homeostasis and protect the respiratory tract (13, 14). Interestingly, despite sharing a common structure throughout the respiratory tract, the cellular composition and response of the airway epithelium varies by anatomical location with nasal and bronchial epithelial cells responding differently (16–18).

To establish colonization, *B. pertussis* employs a diverse array of virulence factors, which include adhesins such as filamentous hemagglutinin (FHA), fimbriae, and pertactin, as well as complement evasion factors and toxins such as adenylate cyclase toxin and pertussis toxin (19). These factors have been shown to disarm the innate immune responses of phagocytic cells, subvert the maturation of dendritic cells and modulate the responses of bronchial epithelial cells (20–23). In addition to these well-characterized virulence factors, *B. pertussis*, similar to its close relative *B. bronchiseptica*, encodes a type III secretion system (T3SS), which injects the cytotoxic effector protein BteA directly into host cells (24–26). While the T3SS in *B. bronchiseptica* is critical for persistent colonization in animal models, including mice, rats and pigs, (27–30), its role in *B. pertussis* pathogenesis is less clear due to a poor expression or absence in laboratory-adapted strains (31, 32). However, clinical isolates of *B. pertussis* have an active T3SS that delivers a less cytotoxic variant of the BteA effector than *B. bronchiseptica*, due to the insertion of A503 into BteA (33). Despite its reduced cytotoxicity, BteA of *B.pertussis* still induces cellular signaling and remodeling of endoplasmic reticulum and mitochondrial network in HeLa cells (34).

In this work, we investigated the responses of human nasal epithelial cells (hNECs), which were cultured from the nasal airway mucosa of healthy donors, to *B. pertussis* strain B1917, a European clinical isolate previously used in controlled human infection studies by the PERISCOPE consortium (35). As the initial site of interaction for *B. pertussis* during respiratory tract infection, hNECs represent a physiologically relevant model. Despite their relevance, they have been underutilized in experimental studies compared to primary bronchial epithelial cells and immortalized bronchial cell lines (23, 36). To the best of our knowledge, there is currently no comprehensive analysis of their responses. We have also investigated the role of the effector protein BteA in modulating the cellular responses of hNECs.

## Results

### Characterization of hNECs cultured at an air-liquid interface

Primary nasal and bronchial epithelial cells grown at the air-liquid interface (ALI) are widely used *in vitro* models for mimicking human airway epithelium (37, 38). However, the utility of these models depends on careful handling of the cultures and differentiation of the cells. To assess the suitability of our hNEC cultures for studying interactions with *B. pertussis*, we first characterized the composition of the generated hNECs.

Primary human nasal cells were isolated from nasal brushings of healthy human donors and expanded by conditional reprogramming as previously described (37, 39). Upon expansion, cells were seeded onto Transwell membranes and air-lifted for 28 to 30 days to promote differentiation. Differentiation into the major airway epithelial subsets was then assessed using multicolor flow cytometry with a sequential gating strategy according to Bonser *et al*, 2021 (40). This analysis included the basal cell marker CD271 (NGFR), the secretory cell marker CD66c (CEACAM5), and the Tubulin Tracker (Tub), which identifies ciliated cells by their high content of acetylated α-tubulin in the cilia. As shown in Fig. 1A, distinct epithelial subsets were identified following gating of single cells. Next, we examined the reproducibility of differentiation and donor-to-donor variability by quantifying subset proportions across three replicates for each of the three donors. Two major cell subpopulations were distinguishable among the single cells – CD66c^+^CD271^-^ and CD66c^-^CD271^+^ subpopulations. Almost all CD66c^-^CD271^+^ cells showed a low Tub signal, indicating that these cells are likely basal cells (40). The CD66c^+^CD271^-^ cells were divided into three subpopulations – CD66c^High^Tub^Low^, CD66c^High^Tub^Mid^ and CD66c^Low^Tub^High^. As shown in Fig. 1B, cell-type proportions were mostly consistent among replicates but displayed inter-donor variability, likely reflecting individual biological differences. The CD66c^High^Tub^Low^ and CD66c^High^Tub^Mid^ cells probably correspond to secretory cells, whereas the CD66c^Low^Tub^High^ cells are likely ciliated cells (40). In order to confirm the accuracy of this classification, we performed fluorescence-activated cell sorting (FACS) of cell subsets from two donors, followed by transcriptomic profiling (Table S1). The heat map of known marker expression (Fig. 1C) validated the subset identities (14, 40) and confirmed the reliability of the flow cytometry panel as well as the differentiation process. Based on the enrichment of marker genes, it also appears that both CD66c^High^Tub^Low^ and CD66c^High^Tub^Mid^ correspond to secretory cells (14).

**Figure 1.**
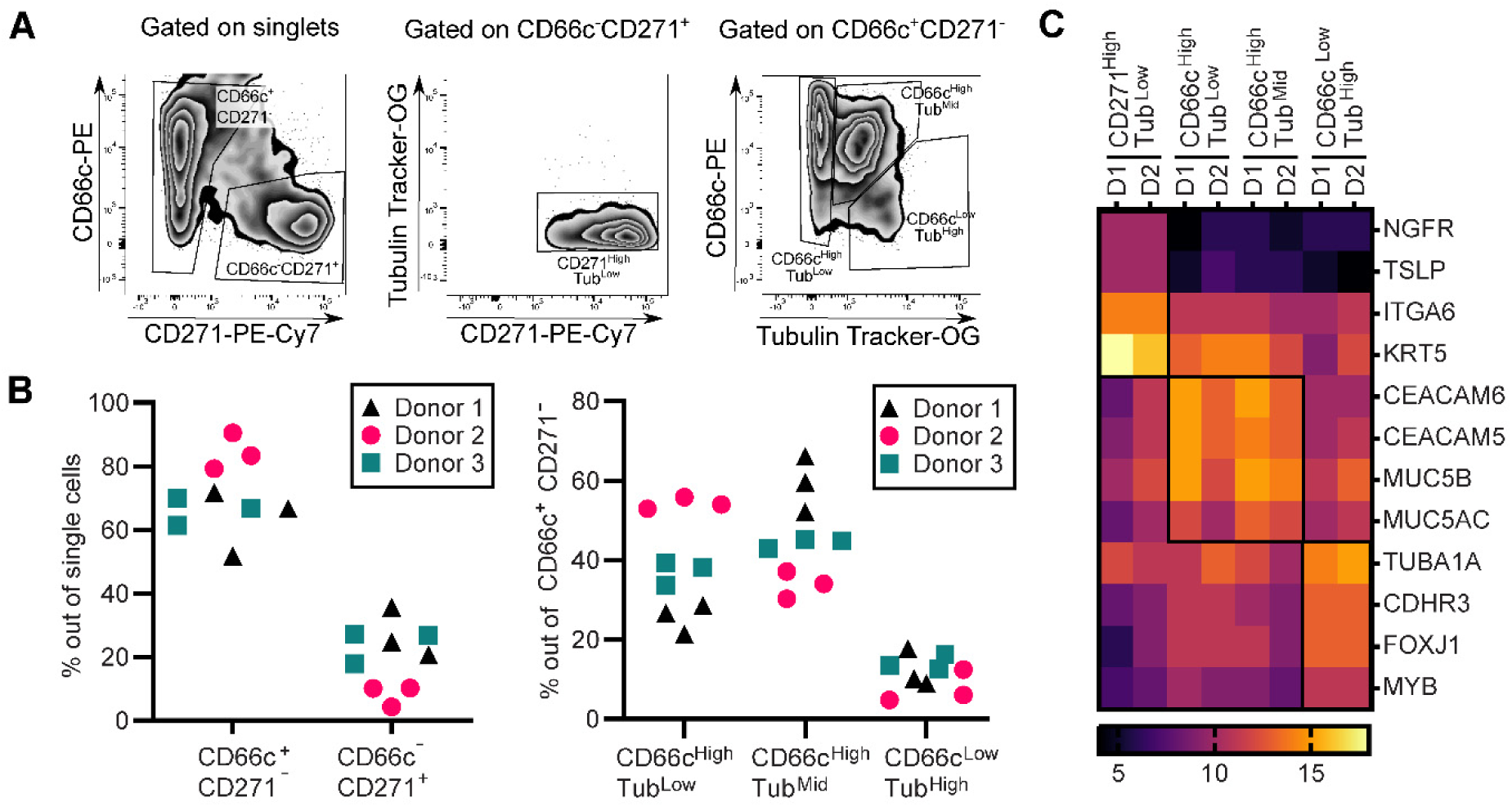
Characterization of hNECs cultured at an air-liquid interface. **(A) Cell gating strategy.** The live single cells were gated according to staining using anti-CD271-PE-Cy7 and anti-CD66c-PE antibodies. The Tubulin Tracker-OG staining was used to refine cell subset identification. Within the CD271 subset, cells with CD271-high/Tub-low expression were classified as basal cells. In the CD66c subset, cells with CD66c-high/Tub-low and CD66c-high/Tub-mid expression were classified as secretory cells, while cells with CD66c-low/Tub-high expression were classified as ciliated cells. These gating strategies were applied to determine donor to donor variability and isolate cell subsets, with specific gates highlighted. **(B) Donor to donor variability.** The percentage of cell subsets was determined in 3 different donors using 3 replicate cultures of hNECs for each donor. **(C) Transcriptome profiling of the cell subsets**. Transcriptome profiling of the isolated cell subsets was performed in two independent experiments to evaluate transcriptional differences. Markers of basal cells: NGFR, TSLP, ITGA6, KRT5, markers of secretory cells: CEACAM6, CEACAM5, MUC5B, MUC5AC, markers of ciliated cells: TUBA1A, CDHR3, FOXJ1 and MYB.

In conclusion, these data demonstrate that our hNEC cultures are differentiated into key epithelial subsets when cultured at ALI and provide a suitable model to analyze the interaction of *B. pertussis* with the nasal epithelium.

### hNECs are permissive for *B. pertussis* infection, with bacteria replicating in mucus and on cell cilia

We then sought to determine the ability of *B. pertussis* to replicate in the hNEC model. For this purpose, we infected cultures of hNECs on Transwell membranes apically with *B. pertussis* strain B1917 in the ALI medium at a multiplicity of infection (MOI) of 10:1. After 6 hours, the medium was carefully removed and the cultures were further incubated without washing under ALI conditions. The bacterial replication was determined from the number of colony forming units (CFUs) grown after bacterial plating at 6, 24, 48, and 72 hours post-infection. CFUs showed a 2-log increase over time reaching 10^8^ cfu per Transwell membrane at 72 hours (Fig. 2A). Interestingly, routine bright-field microscopy during infection revealed no visible cytotoxicity to the epithelial cell layer. Accordingly, as shown in Fig. 2B, infection with an MOI of 10:1 for 24 hours did not result in significant changes in cellular composition, indicating minimal damage in the early stages of infection. To further confirm these data, we visualized ciliated cells, *B. pertussis*, and the integrity of epithelial barrier by fluorescence microscopy. As illustrated in Fig. 2C, within the first 24 hours of infection, *B. pertussis* multiplied mainly in the mucus layer, while multiplication of bacteria on the cell cilia was observed only occasionally at later stages of the infection. This pattern was independent of the donor and also observed in another infection protocol when *B. pertussis* was administered on the top of the hNEC culture on Transwell membrane using five 1-µl drops (see Fig. S1A). As further seen in Fig. 2C and Fig. S1B, staining of the tight junction protein zonula occludens 1 (ZO-1) in the untreated hNECs showed a typical ZO-1 network characteristic of functioning tight junctions and barrier integrity. This network was not affected up to 24 hours following *B. pertussis* infection. However, after 72 hours of infection, the apical ZO-1 network was disrupted and ZO-1 staining was relocated (Fig. 2C and Fig. S1B). Measurements of transepithelial electrical resistance (TEER) also supported these data indicating that the barrier function of *B. pertussis*-infected cells declined at 48 hours, while no change in TEERs was observed at 24 hours post-infection (Fig. 2D). The disruption of barrier function was independent of the T3SS effector BteA, as shown by infections with the mutant *Bp*Δ*bteA* strain, which has an in-frame deletion of the *bteA* gene and a complemented strain *Bp*Δ*bteA::bteA* (Fig. 2D).

**Figure 2.**
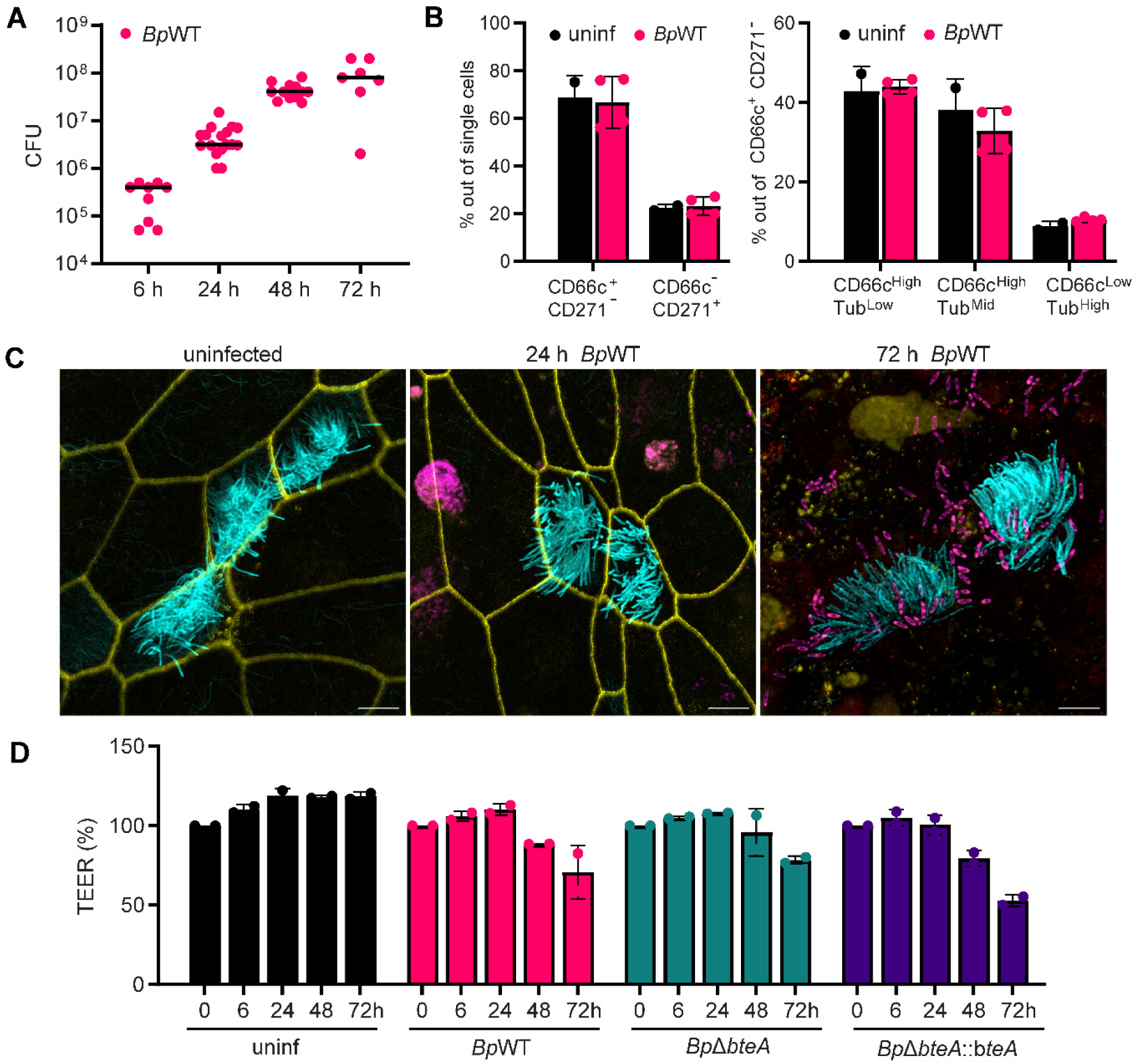
hNECs are permissive for *B. pertussis* infection, with bacteria replicating in mucus and on cell cilia. **(A) hNECs are permissive to infection.** hNECs were apically infected with *Bp*WT at a multiplicity of infection (MOI) of 10:1. After 6 hours, the medium was discarded, and bacterial colony-forming units (CFUs) associated with hNECs per Transwell membrane were quantified immediately or after additional incubation as indicated. Data represent two independent experiments. **(B) Infection with *B. pertussis* does not change the composition of the hNEC culture.** hNECs were apically infected with *Bp*WT at MOI 10:1 and analyzed by flow cytometry 24 hours post-infection as outlined in the legend of Figure 1A. **(C) *B. pertussis* replicates in mucus and on cilia.** hNECs were infected by *Bp*WT expressing fluorescent mScarlet protein. ZO-1 (yellow) was stained using an anti-ZO-1 antibody, followed by an anti-rabbit IgG-DYL405 conjugate, while cilia (cyan) were labeled with an anti-acetylated tubulin antibody, followed by an anti-mouse IgG-AF488 conjugate. Bacteria were visualized by mScarlet expression (magenta). Maximum intensity projections (Z-max) from confocal Z-stack images are shown, and they are representative of two independent experiments. Scale bars, 5 μm. **(D) Transepithelial electrical resistance after infection.** hNECs were apically infected as above with *Bp*WT, *Bp*Δ*bteA,* or *Bp*Δ*bteA::bteA* at MOI 10:1. Transepithelial electrical resistance across the cell layers was measured at the indicated time points in two replicates.

Overall, these results demonstrate that hNEC cultures are susceptible to *B. pertussis* infection and that *B. pertussis* localizes in the mucus layer and only occasionally adheres to the cilia while still continuing to grow in the mucus. The early stages of infection have no effect on the barrier function of the epithelium, which is only impaired in later stages, independently of the BteA effector.

### *B. pertussis* infection triggers a moderate transcriptomic response in hNECs characterized by an increase in mucin production and the absence of inflammatory signaling

To investigate the response of hNECs to infection with *B. pertussis* and determine whether the BteA effector plays a role in the process, we next sought to analyze transcriptomic changes of hNECs 24 hours post-infection. This time point was chosen as it represents the early stage of infection with minimal epithelial disruption and pathology. hNEC cultures generated from the nasal brushings of a single donor (donor 2, Fig. 1B) were infected apically with the indicated strains at an MOI of 10:1, while uninfected controls were treated with medium only. After 6 hours, the medium was carefully removed, and hNECs were further incubated without washing for an additional 18 hours before mRNA isolation, library preparation, sequencing, read mapping, and data analysis.

As shown in Fig. 3A, principal component analysis (PCA) of the transcriptomic data revealed two main clusters for all samples. The first cluster included the data from the uninfected samples, while the second cluster included the data from the infected samples, indicating differences between these two groups. Interestingly, the transcriptomic data of the uninfected samples were quite diverse, probably reflecting the heterogeneity of the individual Transwell membranes. In contrast, data from infected samples clustered more tightly, suggesting minimal differences between responses of hNECs to the different bacterial strains.

**Figure 3.**
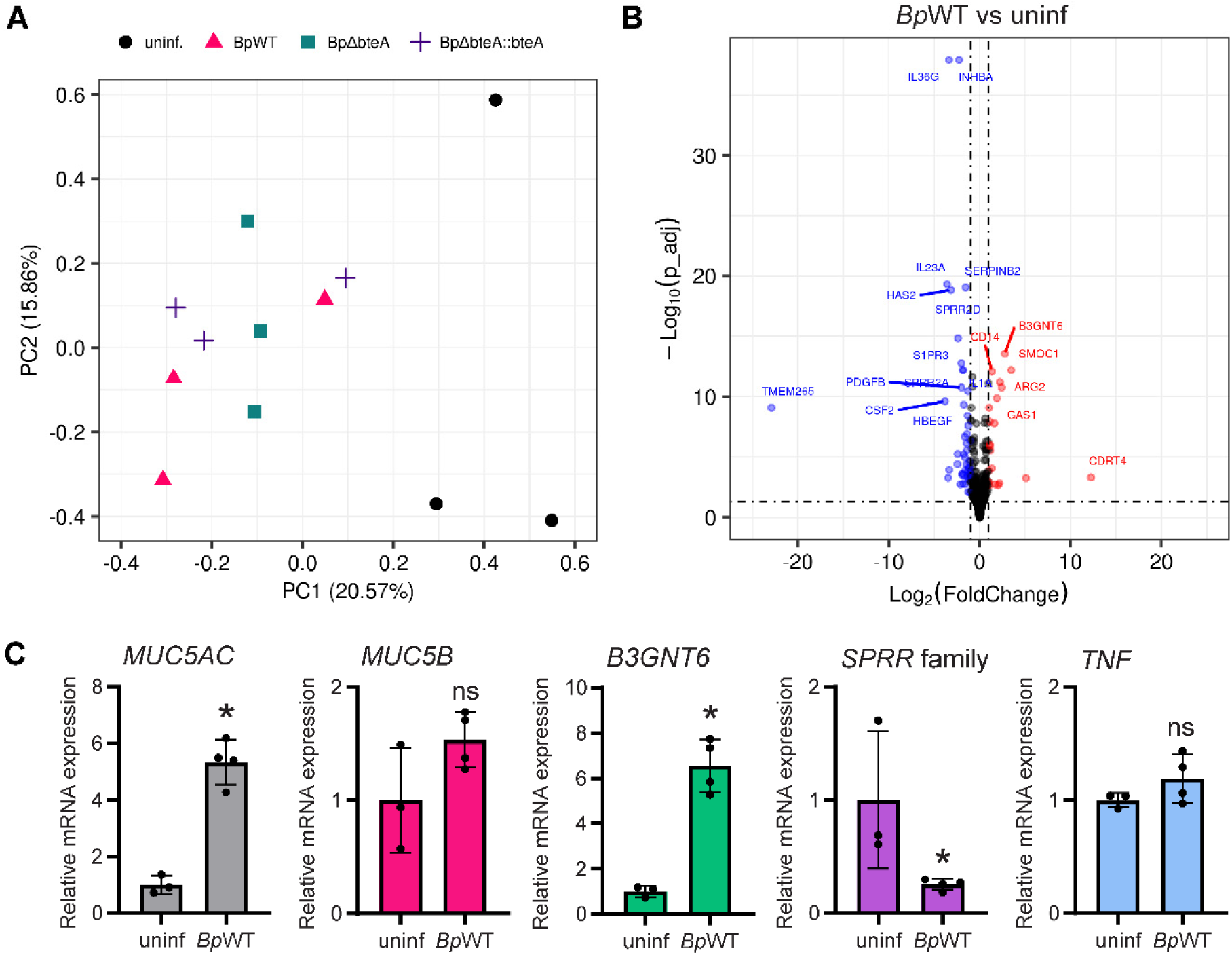
*B. pertussis* induces a moderate transcriptomic response in hNECs characterized by an increase in mucin production and minimal inflammatory signaling. **(A) Principal Component Analysis (PCA) of transcriptomic data.** PCA was performed on transcriptomic data from uninfected hNECs and hNECs infected with *Bp*WT, *Bp*Δ*bteA*, and *Bp*Δ*bteA*::*bteA* at MOI 10:1 for 24 hours. Each dot represents an independent biological replicate. **(B) Volcano plot of differential gene expression in *BpWT-*infected hNECs versus uninfected hNECs.** Significant changes were defined as |log2 fold change| ≥ 1 and adjusted p-value ≤ 0.05. Red dots represent significantly upregulated genes, and blue dots significantly downregulated genes. Black dots indicate genes with nonsignificant changes. **(C) Validation of RNA-seq results by quantitative qRT-PCR.** Relative mRNA expression levels of MUC5AC, MUC5B, B3GNT6, SPRR family, and TNF (TNF-α) were validated using qRT-PCR in *Bp*WT-infected hNECs versus uninfected controls. Samples were collected 24 hours post-infection, and relative expression changes were quantified. Each dot represents independent hNEC culture.

Differentially expressed (DE) gene analysis identified 69 significantly modulated genes following *Bp*WT infection compared to uninfected controls (│log2FC│ ≥ 1; adjusted p-value < 0.05). The results are listed in the Table S3 and illustrated in Fig. 3B with a volcano plot showing the statistical significance (adjusted p-value < 0.05) against the expression change (log2FC). Among the 69 DE genes, 22 genes were upregulated, and 47 were downregulated following the infection. The upregulated genes included the mucin *MUC5AC*, gene involved in mucin glycosylation *B3GNT6,* and the transcription factor *FOXA3*, which promotes mucus production. In contrast, genes for cytokines and chemokines, including *TNF* (*TNF-α), CSF2, IL-1α, IL-1β, IL23A,* and family of the small proline-rich proteins (*SPRR*), possibly exhibiting bactericidal activity (41), were downregulated. These changes were observed in all infected samples compared to uninfected ones, regardless of strain, including *Bp*Δ*bteA* and the complemented strain *Bp*Δ*bteA*::*bteA* (Table S4-S6) and Fig. S2A-B). Validation by RT-qPCR (Fig. 3C) confirmed significant upregulation of *B3GNT6* and *MUC5AC* genes and downregulation of the *SPRR* gene family following *Bp*WT infection at 24 hours post-infection. In contrast, changes in the expression of *MUC5B* and *TNF* were not statistically significant.

To further determine the role of BteA, we also performed DE gene analysis on the transcriptomic profiles of hNECs infected with the *Bp*Δ*bteA* strain compared to *Bp*WT and the complemented strain *Bp*Δ*bteA*::*bteA*. As shown in Table S7-S9 and Fig. S2C-D, only 9 genes reached statistical significance when *Bp*Δ*bteA* strain was compared to *Bp*WT (│log2FC│ ≥ 1; adjusted p-value < 0.05). Of these, 6 genes were significantly downregulated, whereas 3 genes were significantly upregulated. However, only one of these genes, CREB-regulated transcription coactivator 2 (CRTC2), which was downregulated following *Bp*Δ*bteA* infection, showed restored expression upon infection with a complemented strain *Bp*Δ*bteA*::*bteA.* These results are in accordance with PCA, where deletion of the effector BteA had a minimal impact on the overall host transcriptomic response (Fig. 3A), and suggest that BteA does not play a critical role in modulating the host transcriptome within this infection model.

In summary, *B. pertussis* infection elicits a rather moderate transcriptomic response in hNECs, characterized by the upregulation of genes involved in mucin production and the absence of chemokine and inflammatory cytokine induction. The BteA effector appears to have a minimal impact on the transcriptional response of hNECs in this infection model.

### Proteomic changes in hNECs confirm increased mucin production

To further characterize the *B. pertussis*-induced response in hNECs, we next performed proteome profiling at 48 hours post-infection. hNEC cultures, derived from the nasal brushings of a single donor (donor 3, Fig. 1B), were infected apically with the indicated strains at an MOI of 10:1, as described above. After 6 hours, the medium was carefully removed, and hNECs were maintained under ALI conditions for an additional 42 hours. The proteins were extracted, processed for analysis, and subjected to LC-MS/MS. The acquired data were searched against both *B. pertussis* and *H. sapiens* databases.

Analysis of the *B. pertussis* proteins, listed in Table S10, revealed the presence of *B. pertussis* adhesins, including filamentous hemagglutinin (FHA), fimbriae (Fim 3) and pertactin, as well as the autotransporters BrkA and Vag8. In addition, pertussis toxin and adenylate cyclase toxin were detected, although their presence varied from sample to sample, possibly due to the sensitivity of detection and the average bacterial load of 5x 10^7^ of *B. pertussis* per sample (see Fig. 2A). Some of the components of the T3SS, including translocator protein BopD, were detected in some of the infected samples, but the effector BteA was not detected (Table S10).

PCA of the human proteome profiles (excluding *B. pertussis* proteins), shown in Fig. 4A, revealed the clustering of data from replicates within the same experimental groups. Interestingly, there was no clear separation between the proteomic data of uninfected and *Bp*WT-infected samples. In contrast, the proteomic profiles of samples infected with the complemented strain *Bp*Δ*bteA*::*bteA* showed the clearest separation.

**Figure 4.**
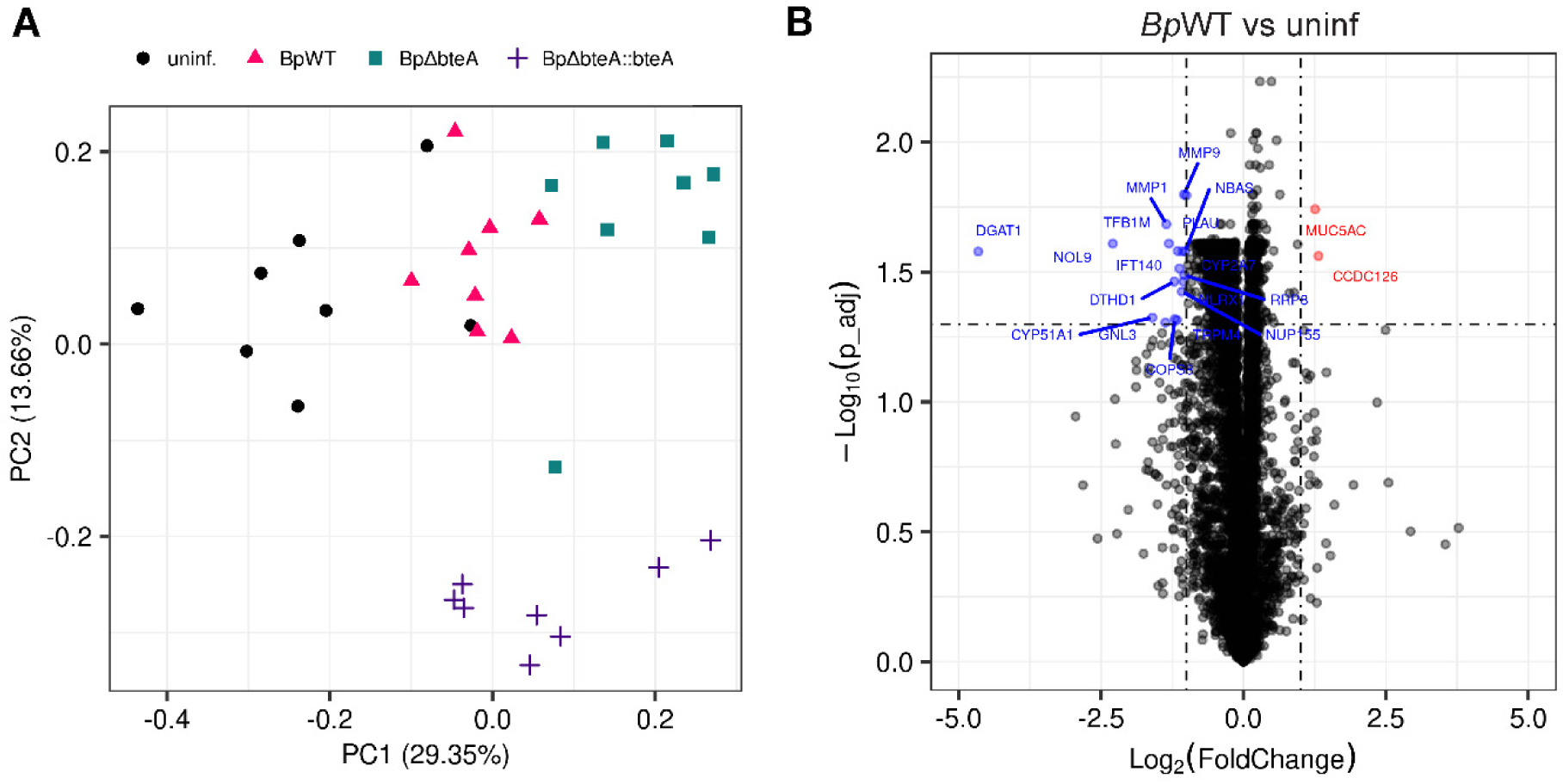
Proteomic changes in hNECs confirm increased mucin production. **(A) Principal Component Analysis (PCA) of proteomic data.** PCA was performed on proteomic data from uninfected hNECs and hNECs infected with *Bp*WT, *Bp*Δ*bteA,* and *Bp*Δ*bteA::bteA* at MOI 10:1 for 48 hours. Each dot represents an independent biological replicate. **(B) Volcano plot of differential protein expression in *BpWT-*infected hNECs versus uninfected hNECs.** Significant changes were defined as |log2 fold change| ≥ 1 and adjusted p-value ≤ 0.05. Red dots represent significantly upregulated proteins, and blue dots significantly downregulated proteins. Black dots indicate proteins with nonsignificant changes.

DE analysis of human proteome profiles revealed significant changes in 20 proteins in *Bp*WT-infected samples compared to uninfected controls (│log2FC│ ≥ 1; adjusted p-value < 0.05), as documented in Table S12 and visualized by the volcano plot in Fig. 4B. Among these, 2 proteins were upregulated, and 18 were downregulated after infection. Importantly, mucin MUC5AC was significantly increased in the *BpWT*-infected samples compared to controls (Fig. 4B). Upregulation of mucins MUC5AC and MUC5B was also observed in samples infected with *Bp*Δ*bteA* and *Bp*Δ*bteA*::*bteA* strains (Table S13-S15 and Fig. S3A-B). Additionally, proteins involved in extracellular matrix remodeling, matrix metalloproteinases 1 and 9 (MMP1, MMP9), and plasminogen activator urokinase (PLAU) were significantly downregulated in all infected samples (Table S15).

To assess the role of BteA, we next performed DE analysis on the proteomic profiles of hNECs infected with the *Bp*Δ*bteA* strain compared to *Bp*WT and the complemented strain *Bp*Δ*bteA*::*bteA*. This analysis, listed in Table S16 and illustrated in Fig. S3C by Volcano plots, identified 26 significantly modulated proteins, of which 5 were upregulated and 21 were downregulated. Among these, only two proteins, tyrosine-protein kinase (BAZ1B) and pleckstrin homology-like domain family B member 2 (PHLDB2), showed restored expression in cells infected with the complemented strain *Bp*Δ*bteA::bteA* (Table S17-S18 and Fig. S3D).

Taken together, these results corroborate the transcriptomic data and confirm that *B. pertussis* infection enhances mucin production in hNECs. Proteomic changes in infected cells, compared to uninfected controls, are moderate and exhibit variability. Proteomic analysis further confirmed that the T3SS effector BteA does not dramatically influence the overall response to *B. pertussis* infection in this hNECs infection model.

## Discussion

This work investigates the interaction of *B. pertussis* with human nasal epithelial cells (hNECs), which are the first cells encountered by *B. pertussis* during respiratory tract infection. We analyzed the dynamics of *B. pertussis* interactions with hNECs, their early host transcriptomic and proteomic responses, and the role of the T3SS effector BteA in these responses. We provide a comprehensive transcriptomic and proteomic dataset for this model. Despite the inherent limitations of the system and variability between donors, we observed increased mucus production and limited inflammatory signaling using both approaches.

In hNEC infection model, *B. pertussis* B1917 localized and multiplied within the mucus layer with minimal attachment to cell cilia up to 24 hours post-infection. This observation contrasts with previous studies, which reported rapid adherence of *B. bronchiseptica* RB50 to cilia in the rabbit tracheal explant model and hNECs (39, 42), as well as attachment of *B. pertussis* 536 and Tohama I strains to the cilia of primary human bronchial cells within 1 and 6 hours of co-incubation, respectively (43, 44). These differences could result from the amount of mucus in the experimental system, the ability of *B. pertussis* B1917 to penetrate the mucus layer, and potential differences in cilia properties and mucus composition. In addition, allelic variations or differences in adhesin expression in *B. pertussis* B1917 and other *B. pertussis* strains, including fimbriae (Fim2 or Fim3), filamentous hemagglutinin (FHA) and pertactin could contribute to the reduced attachment (45, 46). Nevertheless, bronchial and nasal mucus composition have been reported to be highly similar, with only subtle differences (47). Moreover, our proteomic data (Table S10) confirm the expression of Fim3, FHA and pertactin by *B. pertussis* B1917. However, changes in their expression levels or contributions of different alleles compared to the above-mentioned *Bordetella* strains cannot be ruled out.

Despite these differences in the attachment to cilia, *B. pertussis* colonization of the mucus layer is consistent with previous findings showing high bacterial loads in nasopharyngeal washes from infected baboons (48). Interestingly, the major protein components of mucus are mucin glycoproteins, which terminate mainly with sialic acid (49, 50). Both *B. pertussis* and *B. bronchiseptica* have been shown to possess sialic acid-binding capacity, enabling them to bind mucin (51–53). Furthermore, mucin has been shown to prevent *B. pertussis* attachment to A549 lung carcinoma cells (51).

Replication of *B. pertussis* in mucus may also protect epithelial cells from structural damage and activation. Although *B. pertussis* can invade and survive in respiratory epithelial cells (54–56), the invasion rate in primary nasal and bronchial cells is exceptionally low (57), consistent with its extracellular pathogenesis mode. In our model of nasal epithelia, epithelial integrity was preserved for up to 24 hours, as evidenced by ZO-1 staining and TEER measurements, with delayed disruption observed at later stages. This disruption was independent of the T3SS effector BteA, despite its demonstrated cytotoxicity in HeLa cells (34). A likely factor responsible for the barrier disruption is adenylate cyclase toxin (ACT), which has previously been shown to impair epithelial integrity through cAMP signaling (23, 56).

To further investigate the responses of hNECs to *B. pertussis* infection and the role of BteA, we employed transcriptomic and proteomic analyses. Differentiation of hNECs into basal, secretory, and ciliated cell populations was confirmed by flow cytometry using subset-specific markers. However, the system exhibited a pronounced heterogeneity, as could be observed from the proportions of cell subsets detected in cultures from different donors and variable levels of marker protein enrichment (see Fig. 1B-C). Moreover, the donor-specific variability of individual hNEC cultures grown on Transwell membranes was further reflected in the transcriptomic and proteomic datasets, as shown in the PCA plots, emphasizing the inherent limitations of this model.

Transcriptomic and proteomic analyses revealed a moderate response of infected hNECs to *B. pertussis* infection, with 69 significantly modulated mRNA transcripts at 24 h post-infection and 20 significantly modulated protein levels at 48 h post-infection. Importantly, we observed robust upregulation of mucin-related genes, particularly mucin MUC5AC, which, together with MUC5B, represents the major secreted gel-forming mucin of the airways (58). The upregulation of MUC5AC was confirmed at both transcript and protein levels, demonstrating the robustness of this hNEC response during *B. pertussis* infection. Increased mucus production could aid bacterial clearance via mucociliary escalator. However, it is important to mention that MUC5AC, unlike MUC5B, is not essential for mucociliary clearance. Instead, tethering of MUC5AC to the epithelium has been implicated in mucociliary dysfunction and mucus plugging in asthma (58–60). Furthermore, *B. pertussis* has been shown to impair mucociliary clearance by inducing ciliostasis through tracheal cytotoxin (TCT), a muramyl tetrapeptide derived from the bacterial cell wall (36, 61, 62). This impairment may result in mucus stagnation, enabling the formation of a mucus gel matrix that could serve as a niche for *B. pertussis* colonization, thereby facilitating bacterial replication and persistence.

The observed upregulation of MUC5AC aligns with previous studies demonstrating that ACT-induced cAMP signaling mediates mucin upregulation, as shown in the bronchial epithelial cell line VA10 and a mouse model (23, 63). Other pathways, such as Toll-like receptor (TLR) activation and downstream NF-κB signaling, or indirect mechanisms involving high levels of cytokines such as IL-8 and TNF-α may also upregulate mucin production (64–66). However, the TLR activation and the production of inflammatory cytokines appeared to be limited in the hNEC model used in our study. This observation contrasts with the profound inflammatory responses reported in infected macrophages or bronchial epithelial cell line (53, 67). This lack of responsiveness in hNEC could result from low levels of TLR expression and/or reduced availability of cofactors like MD-2, which are essential for effective TLR activation (65, 68). In addition, the apical-basal polarity of human airway epithelial cells further enables to restrict TLR expression to the basolateral surface, and prevents its excessive activation by the nasal microbiota (65). This restriction is often absent in mammalian epithelial cells grown as submerged monolayers. Supporting this, previous studies using ALI cultures of bronchial epithelial cells exposed to higher multiplicities of *B. pertussis* infection than in our system reported only mild immune activation (36). In an infected host, however, the nasal epithelial cell response would likely be modified by interactions with other immune and nonimmune cells and a wide variety of soluble factors. For example, TLR expression on epithelial cells is upregulated during inflammation or in the presence of the microbial community (65). Supporting this, previous studies have identified IL-1β and IFNγ, present in conditioned media derived from *B. pertussis*-infected macrophage and NK cells, are essential for inducing robust chemokine secretion by bronchial epithelial cells in response to *B. pertussis* (36, 69). Interestingly, in our transcriptomic profiling, we have also observed slight upregulation of CD14, lipopolysaccharide (LPS) transferase, helping the binding of LPS to a complex consisting of TLR4-MD2 (70) at 24 hours post-infection. It is, however, also worth mentioning that instead of typical LPS, *B. pertussis* produces lipooligosaccharide (LOS) with a truncated non-repeating O-chain and only 5 acyl chains on lipid A, which might affect its stimulatory properties (71, 72).

In addition to the recruitment and activation of immune cells through the production of cytokines and chemokines, epithelial cells contribute to host defense also by constitutively producing antimicrobial peptides that can be further upregulated upon pathogen recognition. These include antimicrobial peptides of defensin family and possibly also proteins of small proline-rich family SPPR (41, 73). While we observed a slight inhibition of SPRR production at the transcript level, defensin mRNA levels remained unchanged. Importantly, activity of T3SS of *B. bronchiseptica* has been shown to inhibit β-defensin production through NF-kB inhibition (74, 75). However, in our model system, BteA from *B. pertussis* had only a limited effect, as it neither affected epithelial integrity nor profoundly modulated hNEC responses during infection. Since we have recently shown that *B. pertussis* BteA triggers ER and mitochondrial fragmentation in HeLa cells (34), our current results suggest that the function of BteA may be highly context-dependent. Indeed, the regulation of BvgAS-activated T3SS gene expression of *B. pertussis* is complex and not yet fully understood. The *bscN* locus encoding the *B. pertussis* injectisome consists of at least 4 independent regulons regulated by the sigma factor BtrS, the anti-sigma factor BtrA and the recently identified chaperone BP2265 from the CesT family (76–78). This regulatory network likely adapts T3SS expression to as yet unidentified environmental stimuli. Our proteomic data from infected ALI cultures of hNEC did not confirm BteA expression 48 hours post-infection, and we were unable to detect most injectisome components. This may reflect the limited sensitivity of mass spectrometry or may indicate that expression of BteA and/or the T3SS injectisome has been downregulated or is absent. Consistent with this, we also observed downregulation of the T3SS injectisome tip, which consists of the Bsp22 protein, in *B. bronchiseptica* during hNEC infections (39). Furthermore, *B. pertussis* mainly colonized mucus in our system, which might further limit the translocation of BteA by the T3SS injectisome as its delivery requires direct bacterium-host cell contact. This contrasts with secreted toxins, such as adenylate cyclase toxin or pertussis toxin, which can still interact with nasal epithelial cells when *B. pertussis* remains localized in mucus. Further studies using alternative models are essential to clarify the exact role of BteA and T3SS in the pathogenesis of *B. pertussis*. In addition, identification of the host-derived signals that regulate the expression of the BteA and T3SS injectisome will be crucial.

In summary, this work provides a detailed analysis of the human nasal epithelial cells infection by *B. pertussis*, characterized by a delayed disruption of the epithelium, limited epithelial immune recognition, and the exploitation of mucus as a protective niche. All this may allow *B. pertussis* to evade early host defenses and establish a productive infection in the host respiratory tract.

## Methods

### Bacterial strains, growth conditions, and plasmid complementation

The bacterial strains used in this study are listed in Table S19. *E. coli* strains were cultivated at 37°C in Luria-Bertani (LB) agar or LB broth, with chloramphenicol (15 μg/ml) added to the medium when appropriate. *B. pertussis* B1917 strains were grown on Bordet-Gengou (BG) agar medium (Difco, USA) supplemented with 1% glycerol and 15% defibrinated sheep blood (Lab-MediaServis, Jaromer, Czech Republic) at 37°C and 5% CO_2_. For infections, *B. pertussis* was grown at 37°C to mid-exponential phase (OD_600_ 1.0) in a modified Stainer-Scholte (SSM) medium with reduced L-glutamate (monosodium salt) concentration (11.5 mM, 2.14 g/l) and without FeSO_4_.7H_2_O to maximize T3SS expression (33). For plasmid transfer into *B. pertussis* B1917, bacterial conjugation with *Escherichia coli* strain SM10λ pir was used. Complemented *pertussis* strains were selected on BG agar medium supplemented with 15 μg/ml of chloramphenicol and 100 μg/ml of cephalexin, exploiting the natural resistance of *B. pertussis* to cephalexin. Colonies were then re-streaked onto BG agar with chloramphenicol (15 μg/ml) to maintain the pBBRI plasmid.

### Air-liquid interface (ALI) cultures of human nasal epithelial cells

ALI cultures of primary human nasal epithelial cells (hNECs) were prepared as previously described in (39). Briefly, NECs were collected from the nasal brushings of healthy donors using a cytology brush and seeded onto mitomycin-treated 3T3-J2 fibroblasts in NEC medium to enable conditional reprogramming and expansion. After expansion, 5 × 10⁴ NECs were plated on the apical side of a collagen-coated 6.5-mm Transwell filter (Corning Costar). Following 72 hours, the apical medium was removed (air-lifting), and a differentiation ALI medium was used in the basolateral chamber. The ALI medium was replaced three times per week. The air-lifting defined day 0 of the ALI culture, and the experiments were conducted between days 28 to 30. To prevent mucus accumulation, cells were washed with PBS on the apical side every 7 days, starting from days 14 to 17, depending on the mucus buildup as assessed by visual inspection. Differentiation and cell integrity were assessed by visual inspection and transepithelial electrical resistance (TEER) measurement. On day 25, the basolateral medium was replaced with an antibiotic-free ALI medium, with a minimum of two medium changes before experiments.

### Multi-color flow cytometry of hNECs

Multi-color flow cytometry was used to characterize differentiated hNECs. Prior to processing for flow cytometry, hNECs cultured on Transwell membranes were stained with 2x concentrated Tubulin Tracker Green (Invitrogen, Cat. #T34078), applied apically in 200 µl of HBSS. The cells were incubated for 1 hour at 37°C in a 5% CO₂ atmosphere. After incubation, the solution was discarded, and the hNECs were washed twice apically with 200 µl of HBSS. To detach the hNECs from the Transwell membranes, the cells were incubated for 30 minutes in Accutase solution (Invitrogen, Cat. #00-4555-56) supplemented with Collagenase D (1 mg/ml, Roche, Cat. #11088858001) and DNase I (200 µg/ml, Roche, Cat. #10104159001). For this process, 600 µl of the solution was added to the bottom compartment, and 200 µl was applied apically. Following the incubation, the released cells were collected, centrifuged at 350 g for 5 minutes, washed once with flow cytometry buffer (5 % fetal calf serum and 2 mM EDTA in PBS) and resuspended in flow cytometry buffer containing anti-CD66c (clone B6.2, Exbio, Cat. #. 1P-863-T100) and anti-CD271 (clone NGFR5, Exbio, Cat. #. T7-642-T100). Antibody staining was performed on ice for 1 hour. Positive staining was determined using fluorescence-minus-one (FMO) controls to ensure accurate gating and analysis. Data were acquired using BD LSRII flow cytometer and analyzed using FlowJo v10 software.

### RNA sequencing of flow cytometry sorted populations

The cells were stained for multi-color flow cytometry of hNECs as described above, which was followed by sorting of cells on BD Influx cell sorter. Sorted cells were collected directly into RLT lysis buffer, with at least 2 x 10^4^ cells collected per population. Total RNA was extracted according to the instructions of the RNeasy plus micro kit (Qiagen, Cat# 74134). RNA concentration and quality were assessed using the DS-11 spectrophotometer DeNovix. In addition, RNA integrity was analyzed using an Agilent Bioanalyzer 2100 (Agilent Technologies, USA) (see Fig. S4A). Only samples with a minimum RNA integrity number (RIN) of 7 were included for library preparation.

The libraries were prepared using the SMARTer Stranded Total RNA-Seq Kit v2 (Takara Bio, Cat# 634411), using 2 ng of RNA as input per sample. Sequencing was performed on the Illumina NextSeq 500 platform with 75 bp single-end reads. A total of 60 million reads and 30 million reads per sample were obtained for experiment 1 and experiment 2, respectively. Raw sequencing reads were preprocessed with the nf-core/RNAseq pipeline (version 3.10.1) using the default settings for STAR + Salmon (79, 80). Reads were mapped to the ENSEMBL reference genome and transcriptome (*Homo_sapiens* GRCh38.dna_sm.primary assembly genome and GRCh38.110 transcriptome). Gene counts were normalized in Rstudio (version 4.4.2) using package DESeq2 (version 1.44.0) (81) and then transformed with regularized logarithm (rlog). Subsequently batch effect was removed by package limma (82). Visualization of the heat map was done by GraphPad Prism (version 10.3.0).

The data have been deposited with links to BioProject accession number PRJNA1216212 in the NCBI BioProject database (https://www.ncbi.nlm.nih.gov/bioproject/).

### Infection of hNECs and determination of bacterial colony-forming units

Differentiated ALI cultures of hNECs were infected apically with 3×10^6^ CFU of *B. pertussis* B1917 in 200 μL of ALI medium. After 6 hours, the medium was carefully removed without washing. The cultures were either processed immediately or further incubated under ALI conditions for a total infection time of 24, 48, and 72 hours. To obtain bacteria for CFU plating, 200 µl of Accutase solution (Invitrogen, Cat. #00-4555-56) was added to the apical compartment and 600 µl of the same solution was added to the bottom compartment. The cultures were incubated for 30 minutes. After incubation, the apical suspension was carefully pipetted up and down to release the cells and transferred to a tube containing 200 µl PBS supplemented with 2% TX-100. The Accutase solution from the bottom compartment was then used to wash the apical compartment, and the wash was collected and combined with the apical suspension. The resulting suspension was serially diluted tenfold in PBS and aliquots were plated on BG agar plates. Bacterial colonies were counted after 5 days.

### Immunofluorescent staining and image acquisition

Differentiated ALI cultures of hNECs grown on Transwell membranes were infected apically with 3×10^6^ CFU of *B. pertussis* B1917 expressing mScarlet protein under the constitutive *gro*ES promoter (BpWT / mSc, Table S20). Infections were carried out using 200 μL of ALI medium, which was carefully removed without washing after 6 h of incubation, or using 5 µl of SS medium administered as 1-μl drops. At indicated time points, Transwell membranes were fixed with prewarmed 4% PFA (Santa Cruz) for 20 minutes at RT, followed by washes with PBS (3x 5 min). Further, the hNECs on Transwell membranes were permeabilized with 0.2% Triton-X100 (TX-100) in PBS for 10 minutes at RT. The Transwell membranes were then blocked with 4% BSA in PBS supplemented with 0.05% TX-100 (PBST) for 1 hour at RT. For immunostaining, anti-ZO-1 rabbit antibody (ThermoFisher, Cat# 339100) at a dilution of 1:300, and mouse monoclonal anti-acetylated tubulin antibody (Sigma/Merck, Cat# T6793) at a dilution of 1:500 in 1% BSA-PBS were applied overnight at 4 °C. The next day, the Transwell membranes were washed with PBST (3x 5 min) and incubated with the anti-rabbit IgG-DyLight-405 conjugate (Jackson Immunoresearch, Cat# 111-475-003) at a dilution of 1:300, and the anti-mouse donkey IgG-AF488 conjugate (Jackson Immunoresearch, Cat# 115-546-062) at a dilution of 1:500 in 1% BSA in PBST for 1 hour at RT. After incubation, the Transwell membranes were washed with PBST (3 x 5 min) and briefly rinsed in dH_2_O. The membranes were then dissected from Transwells, placed inside AD-Seal mounting spacers (ADVI, Cat# ADS-12-07100), attached to glass slides, and covered with the AD-Mount-F mounting medium (ADVI, Cat# ADM-001). Finally, the membranes were sealed with coverslips No. 1.5H (Marienfeld, Cat# 0107032).

Confocal images were acquired with Leica STELLARIS 8 equipped with a wide-range (440 – 790 nm) light laser with the pulse picker (WLL PP) and highly sensitive hybrid detectors operated by the LAS X software. The objective HC PL APO 40x/1.25 GLYC CORR CS2, WD 0.35 mm, was used with the type G immersion (Leica, Cat# 11513859). DyLight-405 was excited with a 405 nm DMOD laser, and other dyes with the WLL-PLL laser with the following settings: AF488 (491 nm) and mScarlet (561 nm). Z-stacks were acquired with steps of 0.15 μm, and pixel size was set to meet Nyquist sampling criteria. Lightning Expert in the LasX software was used for the deconvolution. Deconvolved images were inspected and processed (brightness/contrast adjustments) with the ImageJ/FIJI software (83), and final figures were prepared in Adobe Illustrator.

### Determination of barrier function by measurements of trans-epithelial electrical resistance (TEER)

To monitor the barrier function of hNECs during differentiation, trans-epithelial electrical resistance (TEER) was measured using Millicell-ERS volt-ohm meter (Millipore, USA). For assessing the barrier function of hNECs following infection with indicated *B. pertussis* strains, TEER measurements were performed using the ECIS Z Theta system (Applied Biophysics) with a plate mounted on the ECIS station. At the indicated time points, 200 µl of PBS was added to the apical side of hNEC cells, and TEER values were recorded. The background resistance of an empty Transwell membrane was subtracted from all measurements. Changes in TEER were calculated relative to the TEER value measured at time point 0 for each individual Transwell membrane.

### Transcriptomic analysis of bulk RNAseq and statistical analysis

At 24 hours post-infection, hNECs on the Transwell membranes were lysed using 0.3 mL of RLT lysis solution (Qiagen RNeasy plus micro kit, Cat# 74134) per membrane. Homogenization was carried out by repeated pipetting to ensure complete cell disruption. Total RNA was then isolated according to the instructions of the RNeasy plus micro kit (Qiagen, Cat# 74134) using single Transwell membrane per sample with the DNAse I treatment. During the RW1 wash step, 80 µl of the RDD buffer containing 10 µl of DNase I solution was pipetted directly onto the spin column, and incubated 15 minutes at RT. RNA concentration and quality were assessed using the DS-11 spectrophotometer DeNovix, and RNA integrity was analyzed using an Agilent Bioanalyzer 2100 (Agilent Technologies, USA) (see Fig. S4B). Samples with a minimum RNA integrity number (RIN) of 6 were used in library preparation.

The libraries were prepared using the KAPA mRNA HyperPrep Kit with polyA selection (Roche, Cat#08098123702), with 300 ng of RNA as input per sample. Sequencing was performed on the NextSeq 2000 Illumina platform with 122 bp single-end reads. At least 25 million reads per sample were obtained. Raw sequencing reads were preprocessed with the nf-core/RNAseq pipeline (version 3.10.1) using the default settings for STAR + Salmon (79, 80). Reads were mapped to the ENSEMBL reference genome and transcriptome (Homo_sapiens GRCh38.dna_sm.primary assembly genome and GRCh38.110 transcriptome). Principal component analysis (PCA) and differential expression (DE) gene analysis were performed using package DEseq2 (version 1.44.0) (81) in Rstudio (version 4.4.2). For both analyses, only genes containing at least 10 reads across all samples were included. PCA was plotted using ggplot2 (version 3.5.1) after data transformation with regularized-logarithm (rlog package DEseq2). Adaptive shrinkage (ashr, (84)) was applied to log2 fold changes to stabilize effect size estimates before DE analysis visualization using ggplot2 (version 3.5.1). Only genes with a |log2 fold change| ≥ 1 and adjusted p-value ≤ 0.05 were considered significantly differentially expressed.

The data have been deposited with links to BioProject accession number PRJNA1214286 in the NCBI BioProject database (https://www.ncbi.nlm.nih.gov/bioproject/).

### Quantitative real-time PCR

RNA isolated as described in the transcriptomic analysis was reverse transcribed into cDNA using the High-Capacity cDNA Reverse Transcription Kit (ThermoFisher Scientific, Cat# 4368814). qPCR was performed using EvaGreen reagent (Solid Biodyne, Cat# 08-25-00001) with 20 ng of cDNA used per reaction with data collected using a CFX384 PCR instrument (BioRad). Two housekeeping genes, *RPL13A* and *GAPDH*, were used as reference genes. The sequences of the qPCR primers and their efficiencies are provided in Table S21 and Fig. S5. Data analysis was done using Biorad CFX Maestro software (Biorad, USA).

### Protein extraction and sample preparation for proteomics

At 48 hours post-infection, hNECs on Transwell membranes were lysed with 0.15 ml of preheated (90°C) SDC lysis buffer containing 2 % sodium deoxycholate and 100 mM Tris-HCl, pH 7.4 per membrane. Lysis was carried out for 20 s, with complete cell homogenization achieved by repeated pipetting. The lysates were transferred to tubes and heated at 95 °C for 5 min. To maximize protein recovery, the membranes were washed with an additional 0.15 ml of preheated SDC buffer, and the wash was pooled with initial extraction. Each sample corresponded to a single Transwell membrane.

Lysates were boiled at 95°C for 10 min in 100 mM TEAB (Triethylammonium bicarbonate) containing 2% SDC (sodium deoxycholate), 40mM chloroacetamide, 10mM TCEP (Tris(2-carboxyethyl)phosphine) and further sonicated (Bandelin Sonoplus Mini 20, MS 1.5). Protein concentration was determined using BCA protein assay kit (Thermo) and 10 µg of protein per sample was used for MS sample preparation. The sample volume was then adjusted to 65 µl in total by adding 100mM TEAB containing 2% SDC. Samples were further processed using SP3 beads on the Thermo KingFisher Flex automated Extraction & Purification System in a 96-well plate. Briefly, 65 µl of the sample was added to 65 µl of 100 % ethanol and mixed with the SP3 beads. After binding, the beads were washed three times with 80% ethanol. After washing, samples were digested in 50 mM TEAB at 40 °C with 1 µg of trypsin for two hours, then another 1 µg of trypsin was added and digested overnight. After digestion samples were acidified with TFA to 1% final concentration and peptides were desalted using in-house made stage tips packed with C18 disks (Empore) according to (85).

### LC-MS/MS and Data Analysis

LC separation was performed on the Dionex Ultimate 3000 nano HPLC system online connected with the MS instrument. Samples were loaded onto the trap column (C18 PepMap100, 5 μm particle size, 300 μm x 5 mm, Thermo Scientific) for 2.000 min at 17.500 μl/min. Loading buffer was composed of water, 2% acetonitrile, and 0.1% trifluoroacetic acid. Peptides were eluted with a Mobile phase B gradient from 4.0% to 35.0% B in 64.0 min. Mobile phase buffer A was composed of water and 0.1% formic acid. Mobile phase B was composed of acetonitrile and 0.1% formic acid. Nano reversed-phase column (Aurora Ultimate TS, 25 cm x 75 μm ID, 1.7 μm particle size, Ion Opticks) was used for LC/MS analysis.

The peptide mixture was analyzed on Thermo Scientific Orbitrap Ascend using a data-independent approach. Eluting peptide cations were converted to gas-phase ions by electrospray ionization in Positive mode. The spray voltage was set to 1600 V and the ion transfer tube temperature to 275 °C. MS1 scans of peptide precursors were analyzed in Orbitrap in the range 145-1450 m/z at 60K resolution and with the following settings: RF Lens 60%, maximum injection time 123 ms, AGC target 100. DIA scans were performed in Orbitrap at 30K resolution. The AGC target was set to 1000, and the maximum injection time mode was set to Auto. Precursor mass range 400-1000 m/z was covered by 30 windows 20 Da wide. Activation type was set to HCD with 28% collision energy.

All data were analyzed and quantified with the Spectronaut 19 software (86) using directDIA analysis. Data were searched against the Human database (downloaded from Uniprot.org in March 2024 containing 20 598 entries) and *Bordetella pertussis* (downloaded from Uniprot.org in June 2024 containing 3 258 entries). Enzyme specificity was set as C-terminal to Arg and Lys, also allowing cleavage at proline bonds and a maximum of two missed cleavages. Carbamidomethylation of cysteines was set as fixed modification and N-terminal protein acetylation and methionine oxidation as variable modifications. FDR was set to 1 % for PSM, peptide and protein. Quantification was performed on MS2 level. Precursor PEP cut-off and precursor and protein cut-off was set to 0.01, protein PEP was set to 0.05. Label-Free Quantification Intensity data were exported (Table S10) and their analysis was performed using Perseus (version 1.6.15.0) (87).

To analyze differences in human proteome after *B. pertussis* infection, PCA and DE analysis were employed. Only proteins detected in at least 6 of the 8 biological replicates per experimental group were included in the analysis. The missing values were imputed using sampling from a normal distribution (Table S11). Statistical significance was determined by non-paired Welch’s t-test and p-values were adjusted using the Benjamini and Hochberg method. Only proteins with a |log2 fold change| ≥ 1 and adjusted p-value ≤ 0.05 were considered significantly differentially expressed. All data visualizations were done via Rstudio (version 4.4.2) with package ggplot2 (version 3.5.1). The mass spectrometry proteomics data have been deposited to the ProteomeXchange Consortium via the PRIDE (88) partner repository with the dataset identifier PXD060316.

## Ethics statement

The sampling of the nasal cavity was conducted on healthy donors who provided written informed consent for the use of their cells in research. The study received approval under EK-297/24 of the Ethics Committee of Motol University Hospital, Prague, Czech Republic.

## Acknowledgments

This work was supported by the grant 21-05466S of the Czech Science Foundation (www.gacr.cz), grant Talking microbes - understanding microbial interactions within One Health framework (CZ.02.01.01/00/22_008/0004597) of the Ministry of Education, Youth and Sports of the Czech Republic (www.msmt.cz) and the Lumina Queruntur Fellowship LQ200202001 of the Czech Academy of Sciences to J.K. We would also like to acknowledge the support of the project LM2023053 (Czech National Node to the European Infrastructure for Translational Medicine) from Ministry of Education, Youth and Sports of the Czech Republic.

---

**Figure S1.**
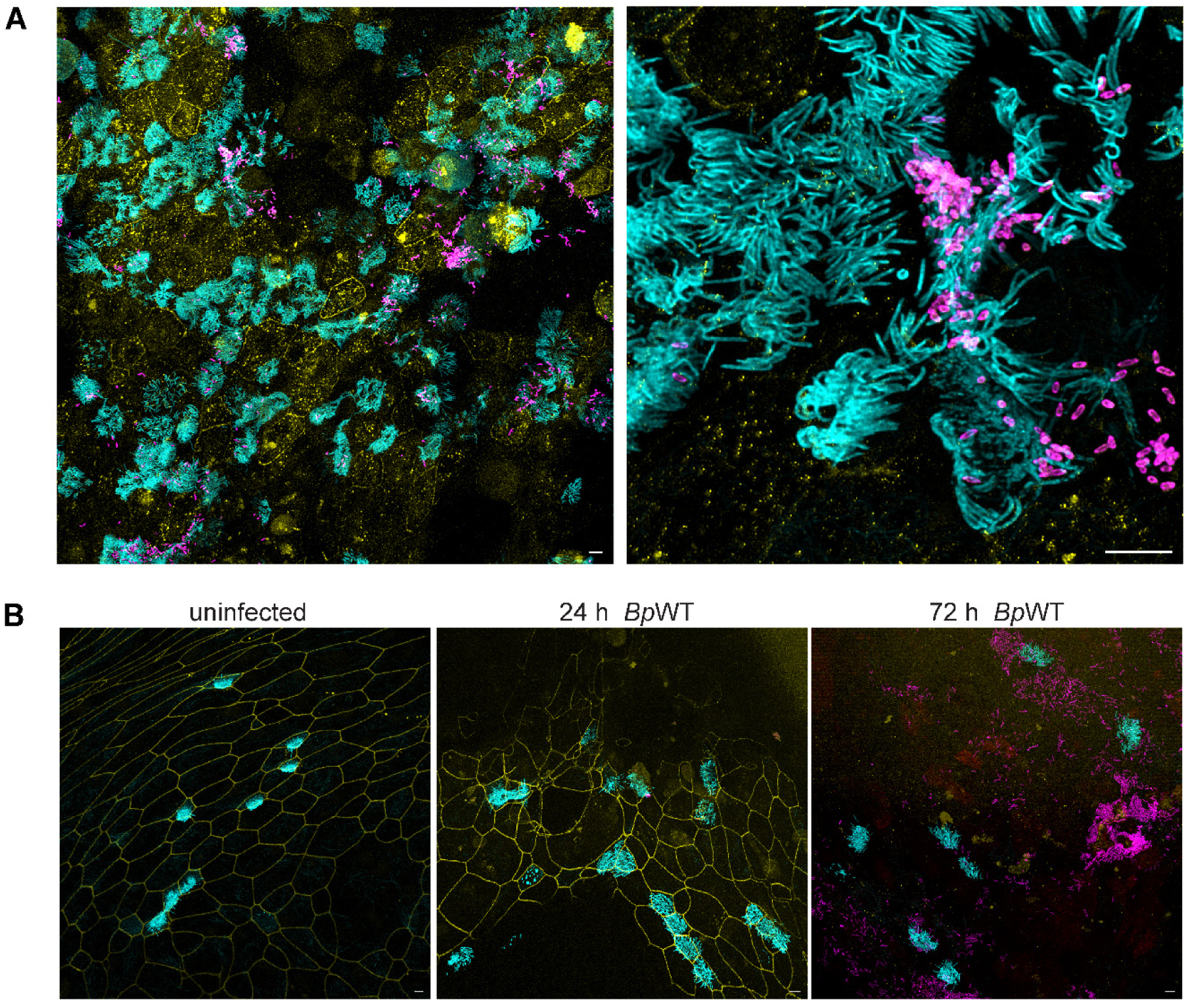
*B. pertussis* replicates in mucus and on cell cilia. *BpWT* expressing fluorescent mScarlet protein at MOI 10:1 was either deposited on the top of the hNECs on Transwell membrane using **(A)** five 1 µl drops or **(B)** 200 μL of ALI medium, which was carefully removed without washing after 6 h of incubation. At indicated time points ZO-1 (yellow) was stained using an anti-ZO-1 antibody, followed by an anti-rabbit IgG-DYL405 conjugate, while cilia (cyan) were labeled with an anti-acetylated tubulin antibody, followed by an anti-mouse IgG-AF488 conjugate. Bacteria were visualized by mScarlet expression (magenta). Shown are maximum intensity projections (Z-max) from confocal Z-stack images. Images are representative of two independent experiments. Scale bars, 5 μm.

**Figure S2.**
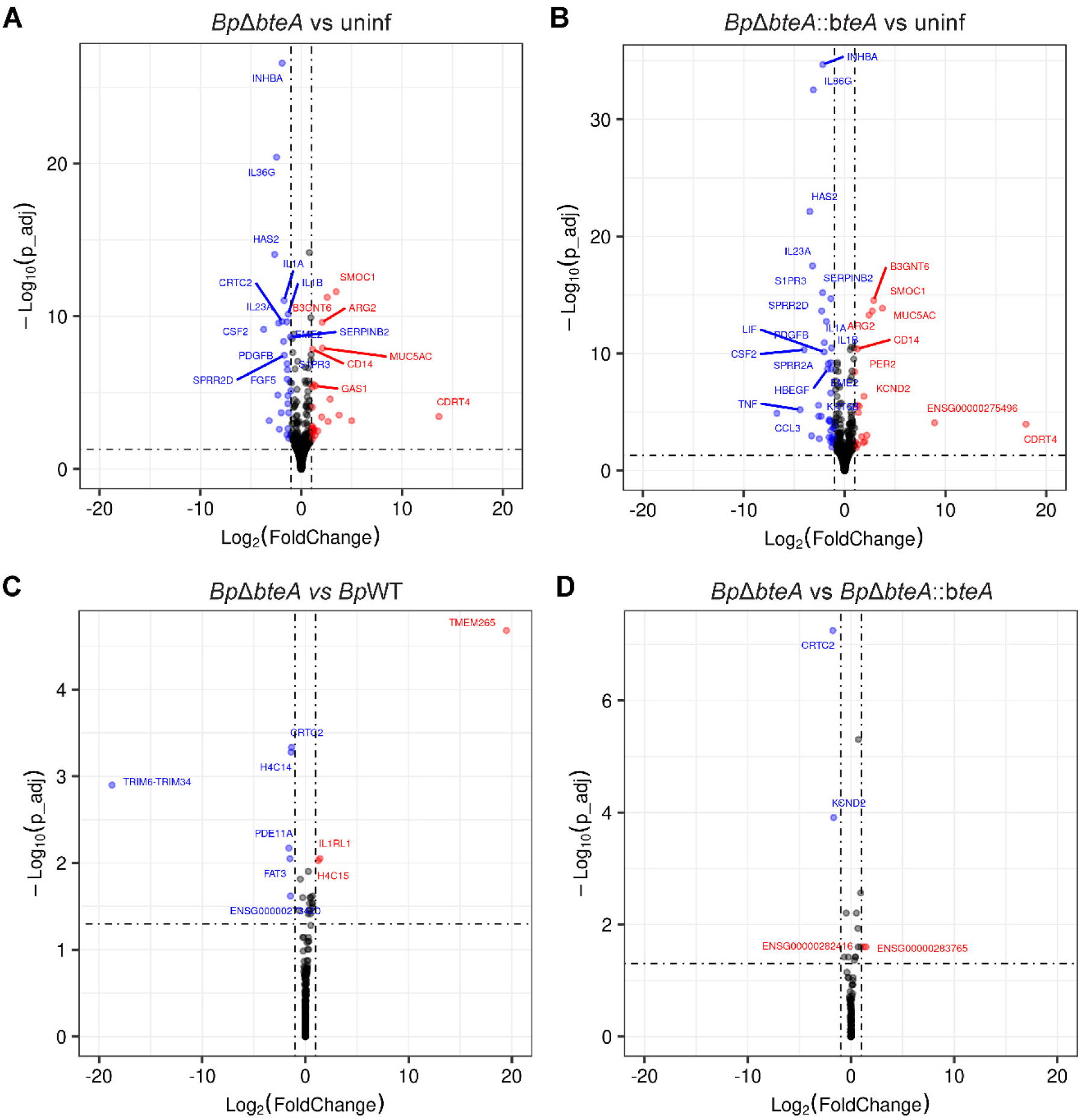
Transcriptomic analysis of hNECs infected by *B. pertussis*. Volcano plot of differential gene expression in hNECs infected with (**A)** *Bp*Δ*bteA* versus uninfected hNEC, (**B)** *Bp*Δ*bteA::bteA* versus uninfected hNEC, **(C)** *Bp*Δ*bteA* versus *Bp*WT and **(D)** *Bp*Δ*bteA* versus *Bp*Δ*bteA::bteA.* Significant changes were defined as |log2 fold change| ≥ 1 and adjusted p-value ≤ 0.05. Red dots represent significantly upregulated genes, and blue dots significantly downregulated genes. Black dots indicate genes with nonsignificant changes.

**Figure S3.**
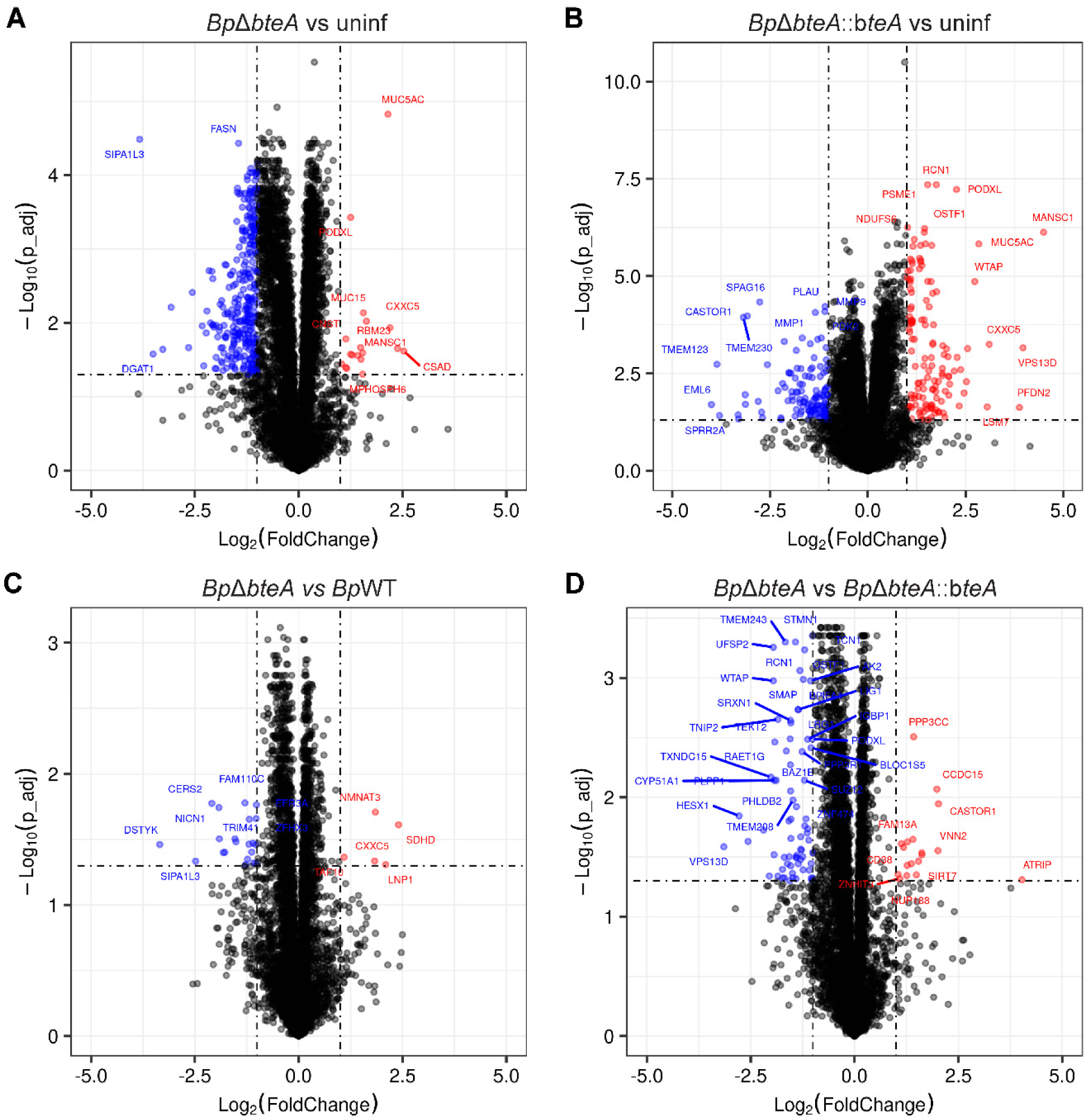
Proteomic analysis of hNECs infected by *B. pertussis*. Volcano plot of differential protein expression in hNECs infected with (**A)** *Bp*Δ*bteA* versus uninfected hNEC, (**B)** *Bp*Δ*bteA::bteA* versus uninfected hNEC, **(C)** *Bp*Δ*bteA* versus *Bp*WT and **(D)** *Bp*Δ*bteA* versus *Bp*Δ*bteA::bteA*. Significant changes were defined as |log2 fold change| ≥ 1 and adjusted p-value ≤ 0.05. Red dots represent significantly upregulated proteins, and blue dots significantly downregulated proteins. Black dots indicate proteins with nonsignificant changes.

**Figure S4.**
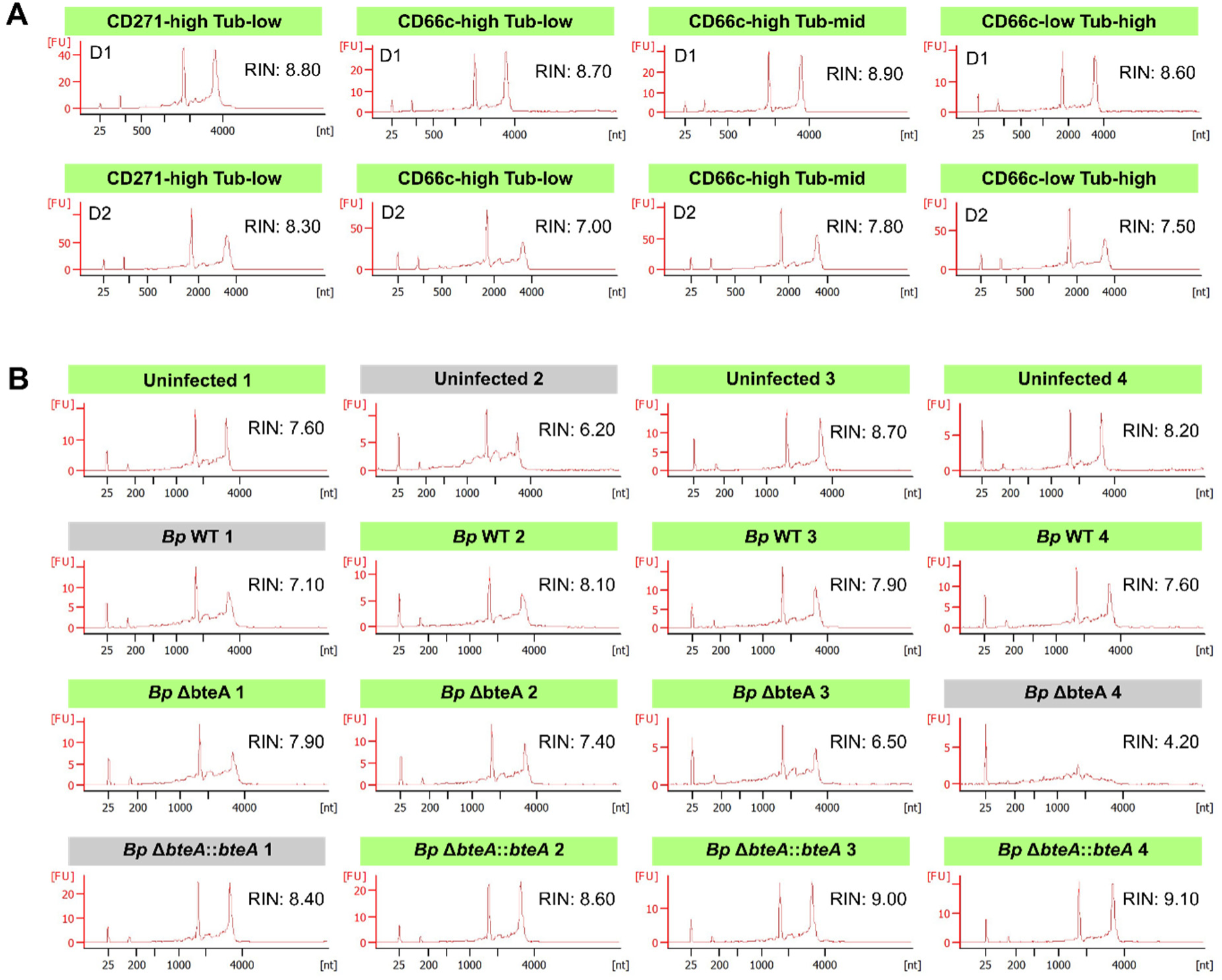
RNA integrity number (RIN) of extracted RNA. The RNA Integrity Number (RIN) of RNA extracted from **(A)** flow cytometry-sorted populations or **(B)** hNEC cultures was assessed using the Agilent Bioanalyzer 2100. Samples highlighted in green were selected for library preparation.

**Figure S5.**
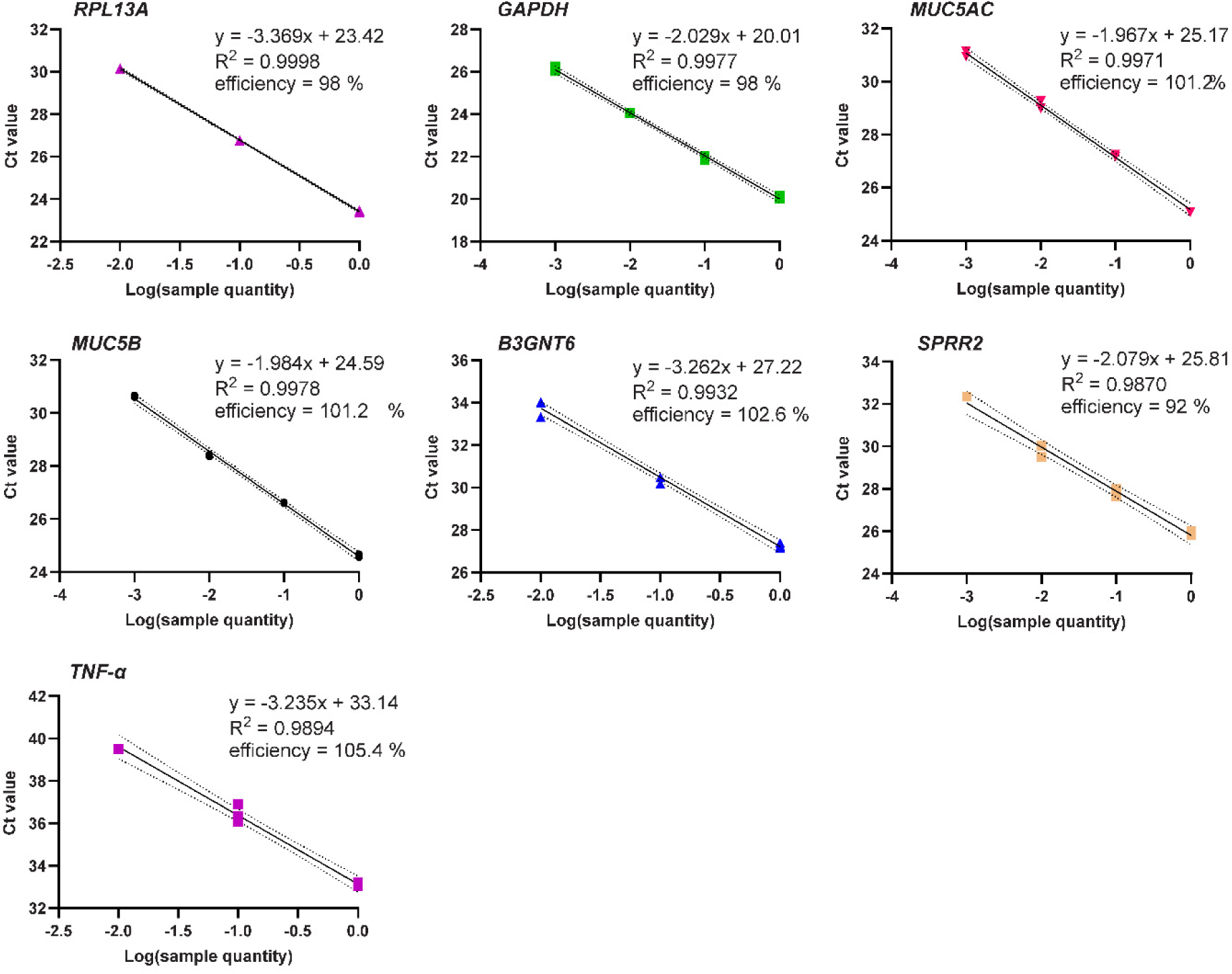
Determination of qRT-PCR primer amplification efficiency. The amplification efficiency of qRT-PCR primers for RPL13A, GAPDH, MUC5AC, MUC5B, B3GNT6, SPRR2 family, and TNF-α was determined using standard curve analysis. Standard curves were generated by plotting the mean CT values against the corresponding serial dilutions of cDNA.

